# Netrin-1 plays a role in the effect of 10 weeks moderate exercise on myocardial fibrosis in rats

**DOI:** 10.1101/348185

**Authors:** Zhou Daliang, Fu Hong, Yu Lifang, Zhang Lingling, Wei Lin, Li Dapeng, Zhang Tianshu, Li Weimin

**Affiliations:** Department of Cardiology, First Hospital of Harbin City, Harbin; Heilongjiang University of Chinese Medicine the First Affiliated Hospital Clinical Pharmacy

**Author notes:** Zhou Daliang, Fu Hong, Yu Lifang contributed equally to the work Li Weimin, Wei Lin.

**Keywords:** netrin-1, acute myocardial infarction, aerobic exercise, rats

## Abstract

This study aimed to determine the effect of Netrin-1 and its receptor on acute myocardial infarction in rats after aerobic exercise.METHODS:Twenty-four rats were randomly divided into three groups: the sham group (n = 8); acute myocardial infarction model group (AMI)(n = 8); and aerobic exercise treatment after acute myocardial infarction group(ET) (n = 8). After 10 weeks, the levels of netrin-1, tumor necrosis factor alpha α(TNF-α), and interleukin 6(IL-6) in the serum were measured. The expression of matrix metalloproteinases 2 and 9(MMP2,9), and their inhibitor, tissue inhibitor of metalloproteinases 2(TIMP2), myocardial netrin-1, Deleted in colorectal cancer(DCC) receptor were evaluated. Histopathological were evaluated. The collagen volume fraction of myocardial tissues was also calculated.RESULTS:Compared to the sham group, the AMI group and ET groups showed increased levels of serum TNF-α, IL-6 and significantly reduced levels of netrin-1. Levels of TNF-α and IL-6 were significantly reduced in the ET group compared to the AMI group, whereas the level of netrin-1 was increased. The expression of myocardial MMP2,9 was significantly increased in the AMI group compared to the sham group, whereas that of myocardial netrin-1, inhibitor of TIMP2 and DCC receptor, was significantly reduced. Compared to AMI group, the ET group showed reduced expression of myocardial MMP2,9 proteins, whereas expression of myocardial netrin-1, inhibitor of TIMP2 and DCC receptor, was significantly increased. The collagen volume fraction of myocardial tissues was significantly increased in the AMI group and ET group compared to the sham group, with the greater increase being noted in the AMI group.CONCLUSIONS: Aerobic exercise could increase levels of serum netrin-1 myocardial netrin-1, and DCC receptor and reduced expression of myocardial MMP2,9 proteins, to improve the degree of fibrosis following myocardial infarction in rats.

## Introduction

Despite modern medical advances, acute myocardial infarction remains one of the main diseases associated with high morbidity and mortality. When myocardial infarction occurs, over the following few weeks, necrotic myocardium is gradually replaced by scar or fibrous tissue, in the natural process of healing. Scarring is conducive to maintaining the normal structure of the heart, and preventing myocardial rupture (1). However, fibrous tissue can also cause the loss of myocardial systolic function, and excessive myocardial fibrosis can induce changes in the shape and function of the heart, a phenomenon referred to as ventricular remodeling. These changes can eventually lead to the development of severe heart failure that can accelerate the death of patients (2–4). Proper control of the degree of myocardial fibrosis is very important to prevent fibrosis of the non-infarcted zone especially.

Aerobic exercise has shown positive results in improving the poor prognosis of patients with myocardial infarction. Studies have shown that aerobic exercise can improve heart function in patients with myocardial infarction, reduce the area of myocardial necrosis, and reduce the degree of ventricular remodeling with low risks [5]. However, many of the associated mechanisms remain unclear.

The role of netrin-1 in cardiovascular disease and the stages of acute inflammation is an emerging area of research. Some studies have shown that an appropriate increase in the concentration of netrin-1, can alleviate myocardial ischemia-reperfusion injury, and reduce atherosclerosis (6, 7). Liu and others (8) reported that aerobic exercise could increase the expression of netrin-1 to relieve cerebral reperfusion injury in rats. However, whether aerobic exercise will show similar effects in myocardial cells has not yet been reported.

Therefore, the present study was undertaken to discuss the netrin- 1 changes after myocardial fibrosis, and evaluate the effect of aerobic exercise on netrin-1 after myocardial infarction.

## Materials and methods

### Animals and groups

Healthy adult male Sprague Dawley (SD) rats, weighing 180–220 g were supplied by the Animal Experiment Center of Harbin Medical University. The rats were randomly divided into the acute myocardial infarction (AMI) model group (n = 8) and the aerobic exercise treatment after acute myocardial infarction group (ET) (n = 8), following the production of an anterior wall myocardial infarction model (described in the following section). Another eight SD rats were allocated to a sham group. All animals were allowed to adapt to the environment in the laboratory over a period of 1 week.

### Myocardial infarction procedure

The AMI and ET groups were subjected to permanent ligation of the left anterior descending coronary artery (LAD), using a 6-0 suture thread to induce myocardial infarction.(9) Under deep anesthesia [ketamine (10 mg/kg body weight)], the thorax was incised at the third intercostal space, and the LAD was ligated 2 mm below the left atrium. Occlusion of the LAD was confirmed microscopically by discoloration of the ischemic area below the ligature. The intercostal space, muscle layers, and skin were closed with continuous sutures using 6-0 silk. Sham animals underwent a similar procedure without occlusion of the LAD.

### Exercise Training Protocol

One week after inducing myocardial infarction, rats of the ET group began to adapt to training on a rat treadmill. The speed was set at 10 to 12 m/min, and rats were initially placed on the treadmill for 10 min every day, for 3 days. After this period of adaptation, the speed was increased to 16 m/min (50%, 60% exhaustion exercise), at a grade of 0°, for 40 min every day, 5 days a week, over 10 weeks(10)

### Exhaustion exercise

Two days after training, the load and movement of the rats were increased. The following protocol was applied: running for approximately 5 m/min, 10 min after warming up, followed by a starting load at 10 m/min, with a gradual increase over 3 min, followed by 5 m/min, until exhaustion.

### Exhaustion judgment standards

The following criteria determined whether the animal was exhausted after the stimulus: inability of animal to keep up with a predetermined speed; hip pressure of rats against the back wall of the cage, while on the treadmill; dragging of the hind legs for 30 s; and behavioral characteristics of deep breathing, large amplitude irregular activity, mental fatigue, and the prone position. Any stimulation of the rats to drive them forward invalidated the determination of exhaustion.

## Determination of IL-6, netrin-1, and TNF-α

Serum interleukin 6(IL-6), netrin-1, and tumor necrosis factor alpha α(TNF-α) were quantified using a commercially available enzyme-linked immunosorbent assay (ELISA) (Cusabio Biotech Wuhan Hi-tech Medical Devices Park, Building B11, #818 Gaoxin Road, Donghu Hi-Tech Development Area, Wuhan, Hubei Province 430206) following the manufacturer’s protocol. Blood was collected via the abdominal aorta, and samples were centrifuged for 10 min at 10,000 *g* at 4°C, and stored at –80°C until further analysis.

## Western blot analysis

The expression of myocardial matrix metalloproteinases 2 and 9 proteins(MMP2,9) inhibitor of metalloproteinases 2(TIMP2), netrin-1 and DCC receptor was measured as follows. Proteins were extracted from all experimental samples and separated by electrophoresis on 8%, 10%, or 12% polyacrylamide gel containing sodium-dodecyl sulfate gel, and then transferred onto a polyvinylidene difluoride membrane. The membranes were incubated for 1 h at room temperature, in 5% nonfat dry milk in Tris-buffered saline solution with Tween 20 (TBST), to block any non-specific binding sites. Membranes were then incubated overnight at 4°C in primary antibodies diluted in a 5% bovine serum albumin solution in TBST. The antibodies used for western blot analysis were as follows: CaSR (1:500), NLRP3 (1:300), ASC (1:200), caspase 1 (1:1000), IL-18 (1:500), IL-1β (1:500), and GAPDH (1:1000). Membranes were subsequently incubated with the secondary antibody for 1 h at room temperature and developed with an enhanced chemiluminescent detection system. The films were scanned and analyzed using the Bio-Rad Image Lab software. All experiments were repeated more than three times.

## Histopathology

After fixation, dehydration, and embedding by paraffin,the myocardial tissue was used to do the hematoxylin and eosin (H&E) and Masson staining Masson staining renders myocardial cells red, and collagen blue. The Motic Med 6.0 software was used for image analysis and measurement of the myocardial collagen volume fraction (CVF). The CVF = myocardial collagen area/test vision area, and values of this parameter were randomly selected from each of six tissue sections. The average of 10 visual measurements from each randomly selected tissue section was recorded.

## Statistical analysis

Statistical analysis of each group was conducted using analysis of variance and the Q test. Statistical analyses were performed with the SPSS 11.0 for Windows software. Statistical significance was set at P < 0.05. Results were reported as mean±S.D.

## Results

### Effect of aerobic exercise on histological assessment of myocardial fibrosis after myocardial infarction

In addition to the sham group, the H&E stained tissues of each group of rats showed varying degrees of inflammatory cell infiltration and cell necrosis, with the most considerable infarction noted in the AMI model group (white arrow, **Fig. 1**). With the exception of the sham group, Masson staining revealed hyperplasia of myocardial cells in the ET group that was significantly less in the AMI group (black arrow, **Fig. 2**). Compared to the sham group, the AMI and ET groups showed significantly higher The collagen volume fraction (CVF) (P < 0.01). However, the CVF of the AMI group showed a more significant increase than that of the ET group (P < 0.05, P < 0.01)(**Fig. 3**).

**Figure 1.**
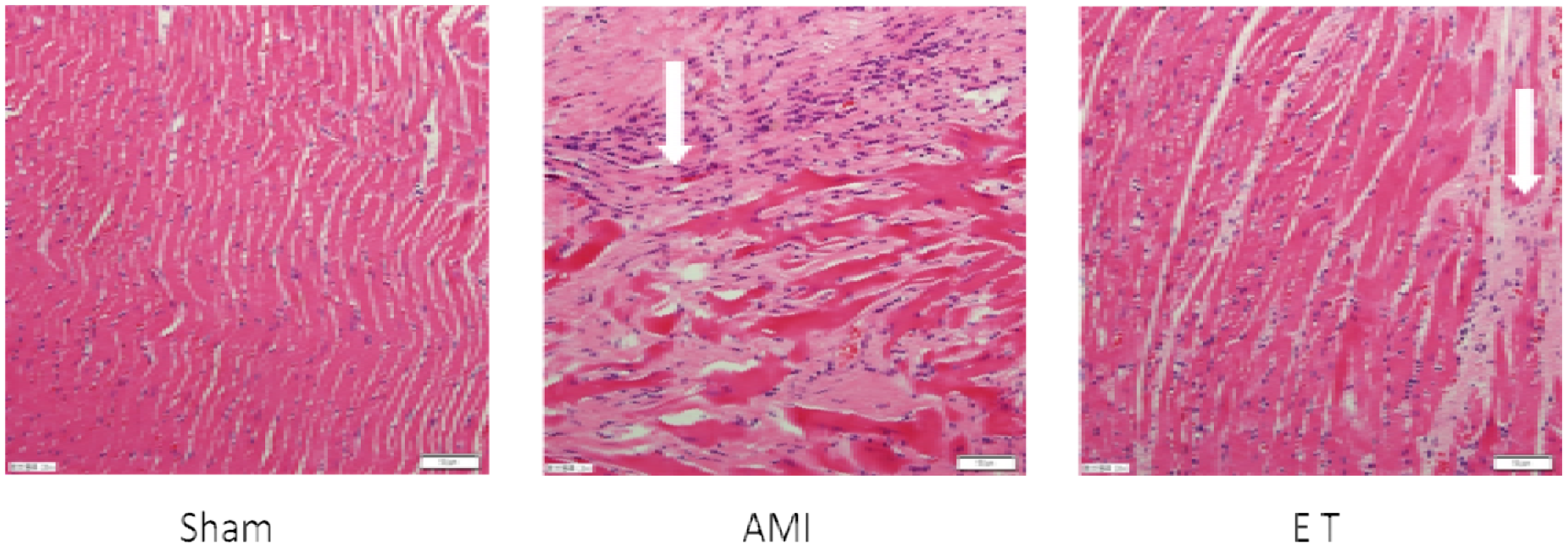
The changes of myocardial tissue structures among groups. Hematoxylin and eosin staining revealed the left ventricles of the myocardium tissues (magnification, ×200).H&E staining of myocardium samples in the three groups showing various degrees of injury. AMI group exhibit the most damage, followed by those in the propofol group.

**Figure 2.**
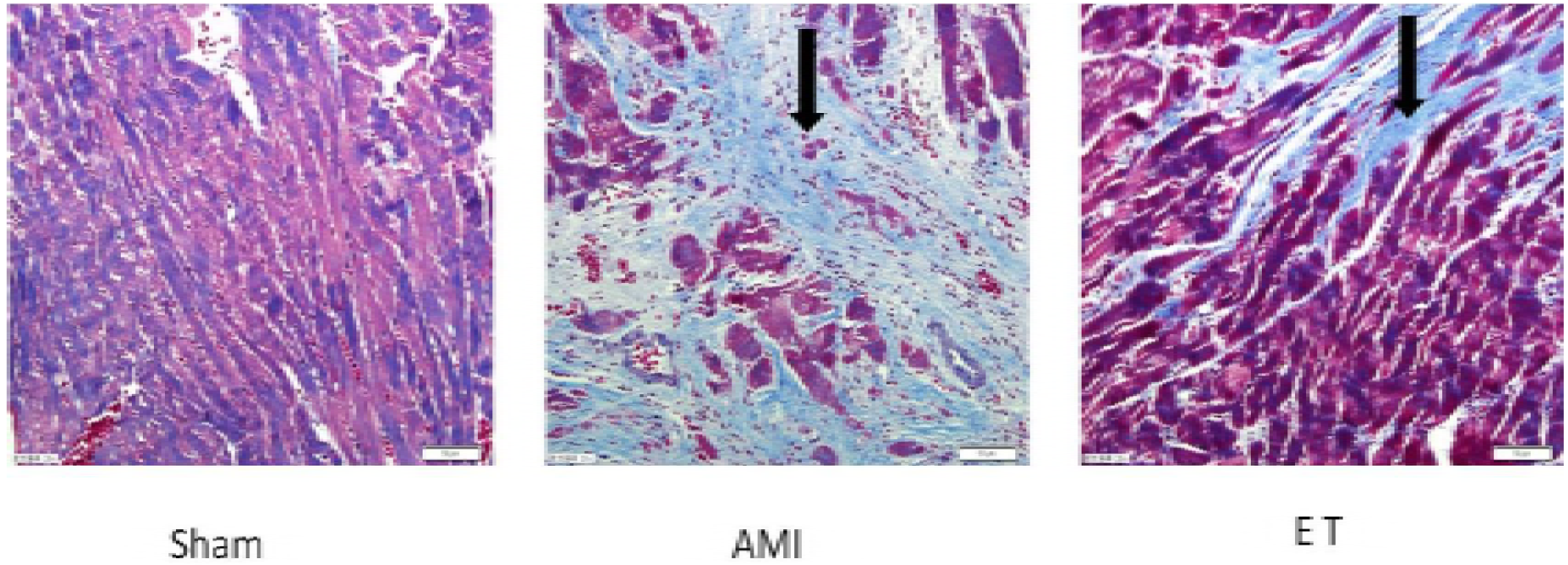
The changes of myocardial tissue structures among groups. Masson staining revealed the left ventricles of the myocardium tissues (magnification, ×200).With the exception of the sham group, Masson staining revealed hyperplasia of myocardial cells in the ET group that was significantly less in the AMI group.

**Figure 3.**
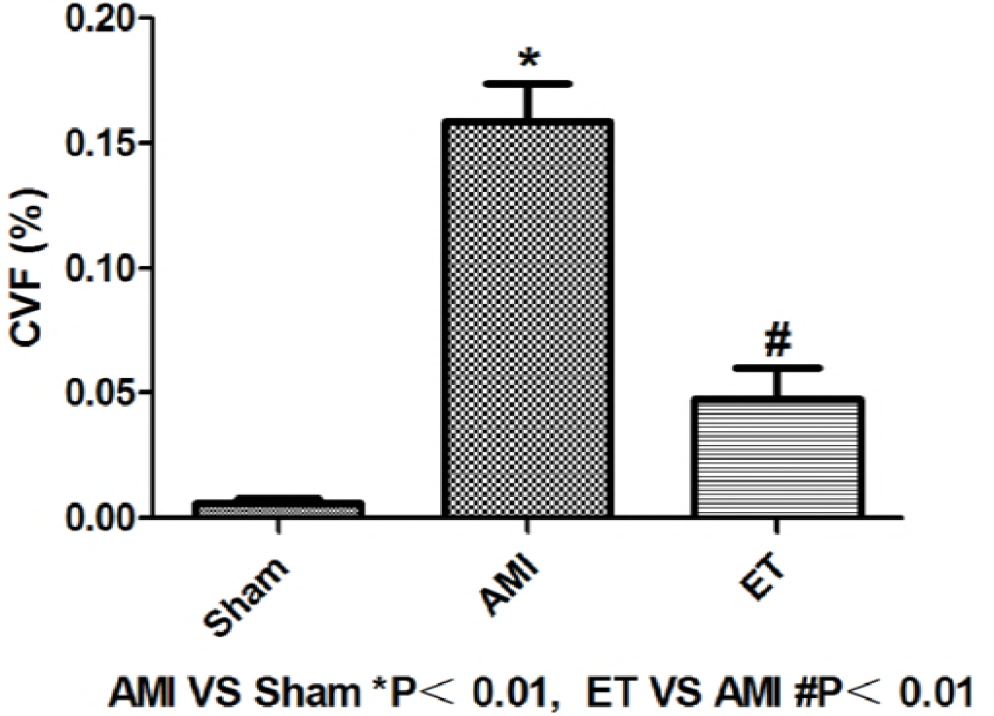
The changes of CVF among groups. CVF, collagen volume fraction; AMI, acute myocardial infarction; ET, aerobic exercise treatment

### Effect of aerobic exercise on IL-6, netrin-1, and TNF-α

Compared to the sham group,in the serum the AMI and ET groups showed significantly higher levels of IL-6, netrin-1, and TNF-α (P < 0.05, P < 0.01). Compared to the AMI group, the ET group showed significantly lower levels of IL-6, netrin-1, and TNF-α(P < 0.05)(**Table 1**).

**Table 1.** Changes in the serum of TNF-α,IL-6,Netrin-1 among groups

| Group | n | TNF-α (pg/mL) | IL-6 (pg/mL) | Netrin-1 (pg/mL) |
| --- | --- | --- | --- | --- |
| Sham | 8 | 18.17 ± 1.12 | 100.48 ± 6.65 | 89.51 ± 6.98 |
| AMI | 8 | 25.89 ± 4.20** | 161.81 ± 14.29** | 73.96 ± 12.67* |
| ET | 8 | 22.16 ± 0.79*▲ | 123.04 ± 28.65*▲▲ | 104.78 ± 31.13*▲ |

AMI, ET vs. Sham \*P < 0.05; \*\* P < 0.01; Sham, ET vs. AMI; ▲ P < 0.05; ▲▲ P < 0.01 AMI, acute myocardial infarction; ET, aerobic exercise treatment.

### Effect of aerobic exercise on protein expression

Compared to the sham group, the AMI groups showed significantly higher expression of myocardial MMP2, MMP9 protein, and reduced expression of myocardial Netrin-1,DCC and TIMP2. Compared to the AMI group, the ET group exhibited significantly reduced expression of MMP2, MMP9(P < 0.05), and significantly increased expression of Netrin-1,DCC and TIMP2 (P< 0.05)(**Fig. 4,5**).

**Figure 4.**
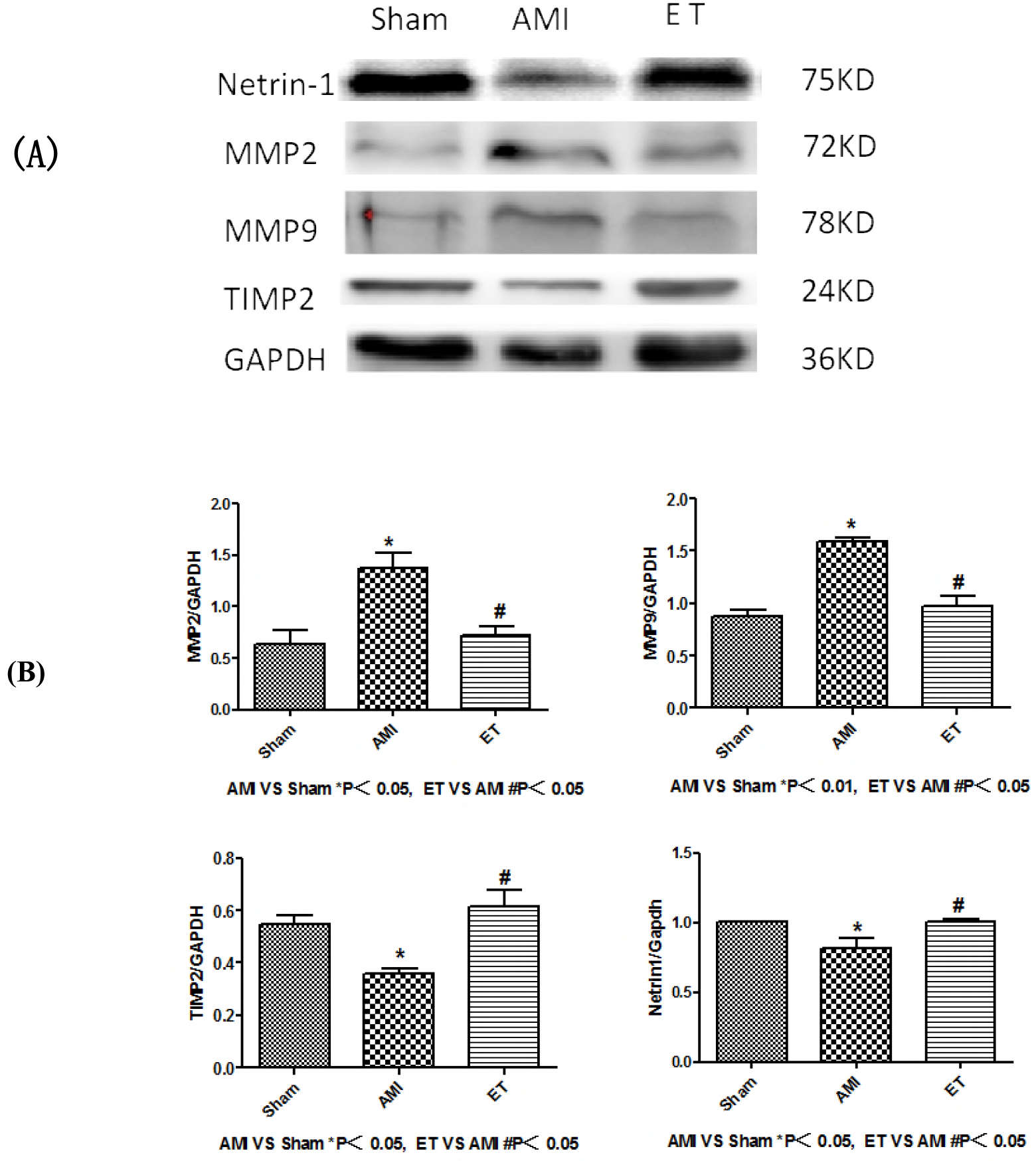
Western Blot analysis of MMP2,MMP9, TIMP2 and netin-1 among all groups.

## Discussion

In this study the H&E stained tissues of rats showed varying degrees of inflammatory cell infiltration and cell necrosis in AMI and ETgroup,but the most considerable infarction noted in the AMI model group, Masson staining also found a large number of blue collagen fibers,in AMI and ETgroup,But the ET group is less. Therefore, this experiment successfully produced the model of myocardial fibrosis rats after myocardial infarction and found that aerobic exercise can reduce the degree of myocardial fibrosis after myocardial infarction.

### Effect of aerobic exercise on the serum netrin-1, myocardial netrin-1 protein and DCC receptors after myocardial infarction

Netrins are a class of laminin-like proteins, which were first identifiedas axonal guidance cues during embryonic development.Netrin-1 is a secreted protein that mediates axonal chemoattractant activity by binding to DCC and neogenin receptors, and mediates repulsion via the uncoordinated-5 (UNC5) receptors (11,12).

The role of netrin-1 in cardiovascular disease and the stages of acute inflammation is an emerging area of research. Some studies have shown that an appropriate increase in the concentration of netrin-1, can alleviate myocardial ischemia-reperfusion injury, and reduce atherosclerosis(6, 7). Studies have also shown that the DCC ERK1 netrin-1/2/eNOS feed-forward mechanism stimulates the aortic endothelial cells to produce NO to protect myocardial cells (13). This a key mechanism, by which regulation of endothelial nitric oxide synthase (eNOS) produces NO to protect myocardial cells in acute myocardial ischemia, chronic congestive heart failure, and ventricular fibrosis (6).Netrin-1 was also reported to control morphogenesisof endothelial cells and vascular smooth-muscle cells.Bouvree et al. demonstrated that netrin-1 inhibits sproutingangiogenesis in cloned chick through UNC5B-binding (14).

In this study, we found that after 10 weeks the AMI group showed reduced levels of serum netrin-1 and reduced expression of netrin-1 protein and DCC receptors in myocardial tissues, whereas the ET group showed significantly increased expression of myocardial netrin-1 protein,DCC receptors and increased levels of serum netrin-1. Therefore, we hypothesized that serum and myocardial netrin-1 expression were decreased after myocardial infarction.But more importantly, aerobic exercise is likely to improve the degree of myocardial fibrosis after myocardial infarction by increasing the expression of serum and cardiomyocardial netrin-1,and its protective mechanism may be in the DCC receptor with netrin-1.

### The effect of aerobic exercise on IL-6 and TNF-α,MMP2, MMP9, and TIMP after myocardial infarction

Myocardial fibrosis after myocardial infarction is an important process of healing. Myocardial infarction that affects a relatively large area following the necrosis of myocardial cells activates inflammatory cells. The role of inflammatory cell necrosis is to remove debris in the myocardial cells and facilitate the reorganization of the extracellular matrix. However, research shows that the levels of a large number of inflammatory cytokines involved in this process, such as IL-6 and TNF-α, become increased, thereby promoting the activation of inflammatory cells (2). Moreover, excessive activation of inflammatory factors disrupts the stability of the extracellular matrix environment, and further promotes the excessive activation of MMPs, thereby accelerating the process of fibrosis after myocardial infarction, and subsequent deterioration of cardiac function (15, 16).

There is considerable collagen deposition following AMI, because of the reconstruction of myocardial cells and the extracellular matrix in the infarcted area. Significant hyperplasia of collagen positive cells serves to prevent myocardial rupture, and is a natural process of self-protection. However, this process can also increase myocardial stiffness in the infarcted area and reduce its compliance, thereby causing cardiac diastolic dysfunction or failure (1, 17). The MMPs are secreted proteins that are produced mainly by neutrophils, macrophages, smooth muscle cells, and endothelial cells. The MMPs play a role both in matrix degradation and in the regulation of collagen synthesis. The inhibitor of MMPs, TIMPs, regulates inhibition under physiological conditions, to maintain a dynamic balance between the actions of MMPs and TIMPs. This effectively coordinates the degradation and reconstruction of the extracellular matrix, and maintains the integrity of its organizational structure and stability of the internal environment (18).

Among the family of MMPs, MMP2 and MMP9 are two important members associated with the modulation of myocardial fibers, and their activity often causes both increased levels of collagen and the aggravation of fibrosis; however, a decline in their activity often leads to reduced fibrosis (19). The present study shows that the activity of MMP2 and MMP9 increases dynamically after myocardial infarction, and this change is positively correlated with cardiac function and ventricular remodeling (20).

In this study, we found that 10 weeks after myocardial infarction in rats, levels of serum inflammatory cytokines IL-6 and TNF-α were both significantly increased. Different levels of collagen deposition were noted in the infarcted area, in response to the inflammatory cell infiltration and the subsequent necrosis of myocardial cells was observed. Thus, the characteristics of myocardial fibrosis also affected the surrounding tissue. in this study confirms that aerobic exercise can reduce the activation of inflammatory factors, enhance the reaction of inflammatory cells, reduce the expression of MMP2 and MMP9 protein in rats, and increase the expression of TIMP-2. Based on measurements of the CVF, we discovered that aerobic exercise could reduce the CVF following myocardial infarction, and reduce the degree of myocardial fibrosis. Thus,this study suggests that aerobic exercise can reduce fibrosis associated with myocardial infarction in the myocardial cells of rats. The mechanism by which this is achieved might be related to the inhibition of inflammatory cytokines, and regulation of the expression of MMP2, MMP9, and TIMP proteins.

In conclusion,the experiment successfully explored the protective effect of aerobic exercise on myocardial infarction, and As far as we know, this is the first study to explore the relationship between aerobic exercise and the expression of myocardial netrin-1,and netrin-1 is likely a novel target for therapeutic intervention, following myocardial infarction and myocardial fibrosis,but many mechanism is not clear, this is also our team to discuss important direction in the future.

### Limitations of the study

All results presented in this study were obtained from the analysis of male mice, and therefore cannot be directly extrapolated to female mice. To our knowledge, no information is currently available regarding potential sex-specific exercise-related differences in post-myocardial infarction remodeling.

**Figure 5.**
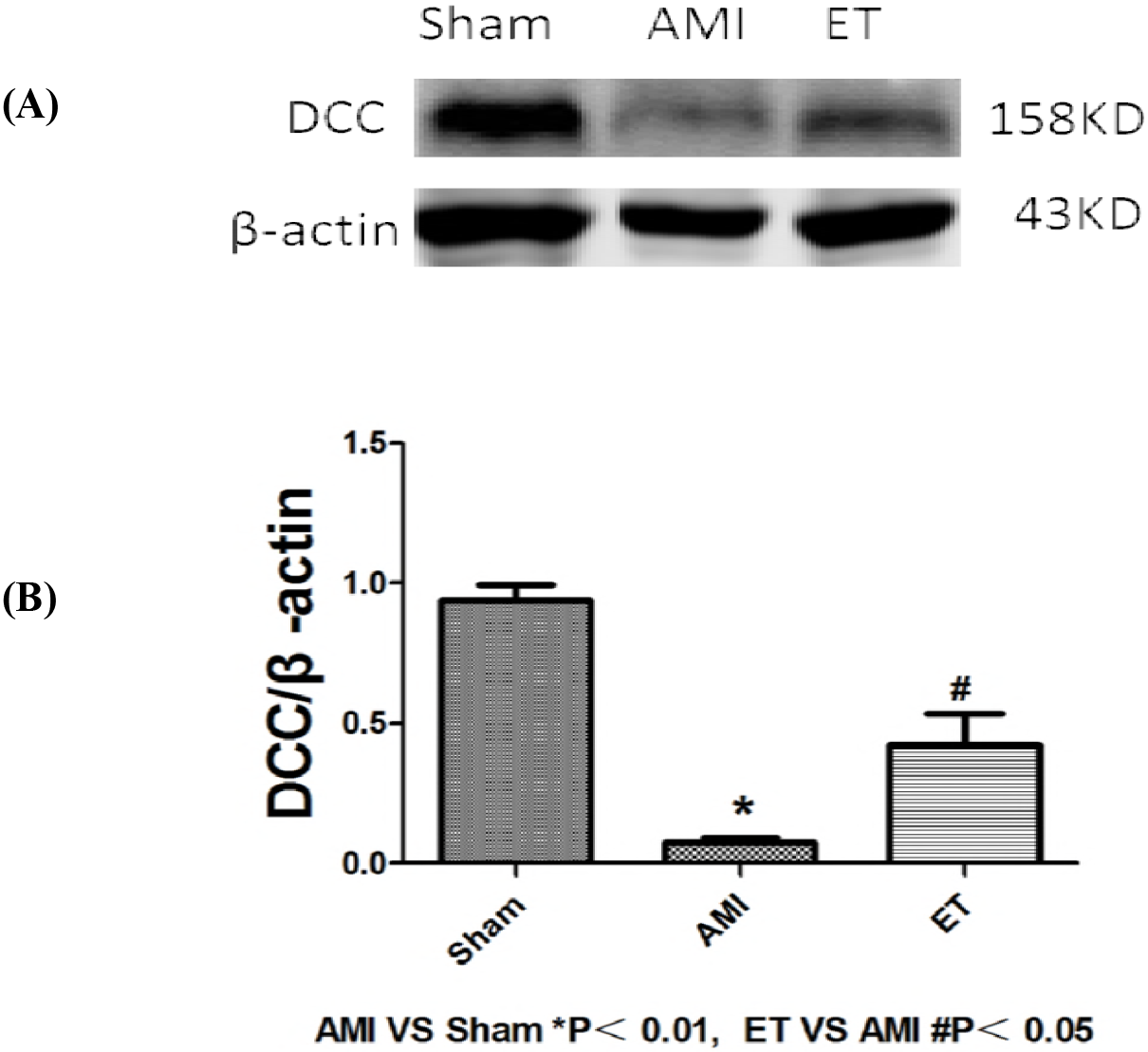
Western Blot analysis of DCC among all groups.

